# First density and abundance estimate for the only East African population of Western bongo (*Tragelaphus eurycerus eurycerus*) in Semuliki National Park, Uganda

**DOI:** 10.64898/2026.09.23.753876

**Authors:** Tommy Sandri, Naomi Matthews, Sam Isoke, Magloire Vyalengerera, Caroline Asiimwe, Robert Aruho, Achaz von Hardenberg, Stuart Nixon

## Abstract

Information on Western bongo (*Tragelaphus eurycerus eurycerus*) across its range remains scarce and fragmented, leaving the status of populations poorly understood, limiting effective conservation planning. We conducted three camera-trap surveys across Semuliki National Park, Uganda, between October 2019 and August 2020 (90 camera locations; 11,640 camera days), during which we detected the only currently known population of Western bongo in East Africa. We used spatially explicit capture–recapture (SECR) models to estimate population density using data from the final survey (March–August 2020), applying an individual identification system adapted from Eastern bongo (*Tragelaphus eurycerus isaaci*). We included sex and habitat covariates derived from remote sensing as predictors of density. Bongos were detected on 17 occasions, and we identified 18 individuals (11 females and 7 males), of which seven immature individuals (5 females and 2 males). The top-ranking SECR model included the tasseled cap greenness (an index of vegetation productivity) and sex as predictors of density. The model estimated a population size of 56.49 individuals (SE = 7.69; 95% CI = 41.43–71.56), with an estimated density of 0.58 individuals km^−2^ (SE = 0.171; 95% CI = 0.32–1.09) within Semuliki National Park. These results provide the first quantitative baseline for this population and suggest the presence of an established breeding population, highlighting the need for structured monitoring to support its long-term conservation.

## INTRODUCTION

The bongo (*Tragelaphus eurycerus*) is the largest forest antelope and occurs across the equatorial forests of Africa (Elkan & Smith, 2013). Two subspecies are recognised: the Western (lowland) bongo (*T. e. eurycerus*), distributed across West and Central Africa, and the critically endangered Eastern (mountain) bongo (*T. e. isaaci*), restricted to the highlands of central Kenya (Faria et al., 2011; Sandri et al., 2023). Camera trap surveys in 2018 in Semuliki National Park, Uganda, (hereafter ‘Semuliki’) documented the first known population of Western bongo in East Africa (Chester Zoo, 2019; Gibbon & Sever, 2022), representing an eastward extension of the species range beyond its known limit in the Ituri forest in eastern DRC (Elkan & Smith, 2013). These findings also suggest that Uganda may be the only country where both subspecies historically occurred, given records of Eastern bongo from Mount Elgon until the early 20th century (Kingdon, 1982). Despite being more widespread and abundant than its eastern counterpart, information on the status of Western bongo remains limited across much of its range (IUCN SSC Antelope Specialist Group, 2017). Establishing baseline population information for the Semuliki population is therefore essential for informing conservation planning. Estimates of population size, density, and sex ratio establish a benchmark against which future population change can be assessed (Callaghan et al., 2024), enabling the evaluation of threats, the effectiveness of management interventions, and the prioritization of conservation actions (Yoccoz et al., 2001).

Forest antelopes are notoriously difficult to study and monitor, but camera trapping has proven effective for investigating their ecology and distribution (Amin et al., 2016; Bowkett et al., 2008; Gray, 2018; Sandri et al., 2023). Bongos can be individually identified by the unique stripe patterns on their flanks (Gibbon et al., 2015; Sandri, 2020; Sandri et al., 2023b), enabling the application of capture–recapture approaches to estimate population density. While traditional capture–recapture models estimate population size (Petit & Valiere, 2006; Otis et al., 1978), the spatial structure of camera trap data allows the use of spatially explicit capture–recapture (SECR) models, which explicitly model the detection process in space and directly estimate density without requiring a separate estimate of the effective sampling area (Efford & Fewster, 2013). Bongos are considered habitat specialists associated with dense understory and secondary vegetation (Estes et al., 2008; Sandri, 2020). Incorporating habitat covariates within SECR models can therefore improve density estimation in structurally heterogeneous forest systems (Kristensen et al., 2018).

Here, we provide the first estimates of density and abundance for Western bongo in Semuliki using SECR models incorporating habitat covariates to account for the species ecology (Green et al., 2020; Estes et al., 2008). These results provide initial insights into the status of this newly documented population and establish a baseline for future monitoring (Martin et al., 2007; Nichols & Williams, 2006), supporting conservation planning for the only known population of this antelope in East Africa.

## METHODS

### Study Area

Semuliki (~220 km^2^), located in southwestern Uganda, represents the easternmost extension of low- and mid-altitude Congo Basin rainforest and contains the only remaining tract of true lowland tropical forest in East Africa (Howard, 1991; Plumptre et al., 2007). The park is contiguous with the northern sector of Virunga National Park in the Democratic Republic of Congo and forms part of the Greater Virunga transboundary landscape within the Albertine Rift (Plumptre et al., 2007). The park lies northwest of the Rwenzori Mountains and spans an altitudinal range of 676–760 m above sea level (Howard, 1991; Figure 1). Vegetation is predominantly moist evergreen to semi-deciduous, dominated by Uganda ironwood (*Cynometra alexandri*), with a mosaic of additional habitats including riverine forest, seasonally inundated swamp forest, papyrus swamps, secondary forest, and small grassland patches (Howard, 1991).

**Figure 1:**
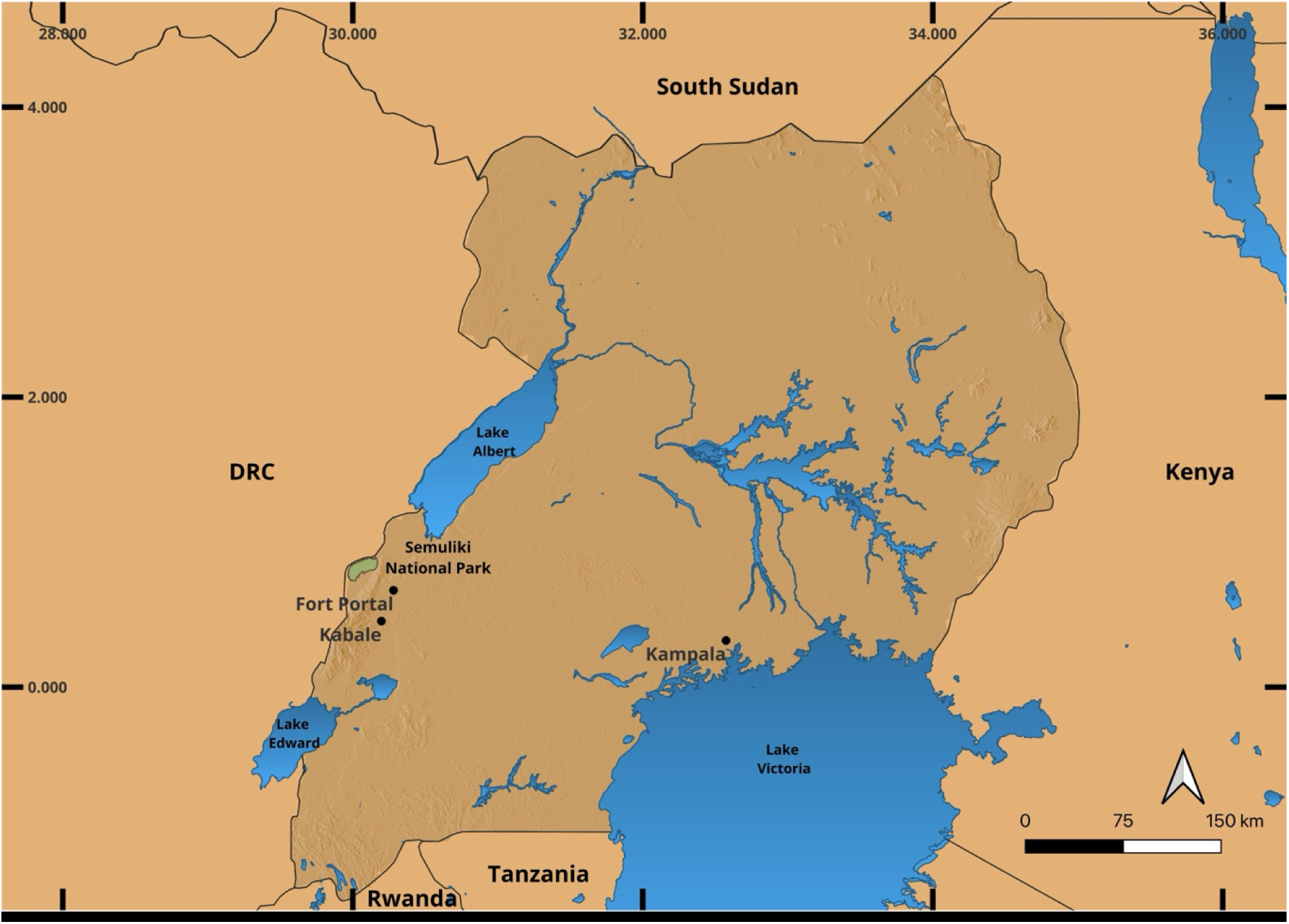
Location of Semuliki National Park within Uganda

Historical anthropogenic disturbance, including periods of settlement and forest exploitation, has contributed to the prevalence of secondary forest within the park. Despite this, the park retains strong ecological affinities with central African forest systems and supports high levels of biodiversity characteristic of the Albertine Rift (Plumptre et al., 2007).

### Camera Trap Survey

Camera trap surveys were conducted between October 2019 and August 2020 as part of a broader wildlife monitoring programme (Matthews et al., 2023), primarily focused on Giant pangolin (*Smutsia gigantea*). Surveys comprised three camera trap deployment arrays: two consecutive arrays in the eastern sector (October–December 2019; December 2019–January 2020) and one array in the western sector (March–August 2020; see Figure 2). Bongos were detected only during the surveys conducted in the western-sector; consequently, all analyses presented here are based exclusively on data from this survey, while the earlier eastern surveys are described for completeness. A 500 × 500 m grid was overlaid on the eastern and western sectors of the park, and grid cells were randomly selected for camera deployment. Within selected cells, cameras were positioned as close as possible to the grid centroid, with placement preferentially along animal trails or areas showing signs of wildlife activity. In the absence of such signs, cameras were placed at the centroid. Detector spacing averaged 843 m (SD = 225 m; median = 796 m), reflecting a broadly regular grid with some variation in camera placement due to logistical constraints of the terrain. Camera traps (Browning Recon Force Advantage and Reconyx HyperFire HC550 and HP2W) operated continuously (24 h day^−1^) and were configured to record videos when motion was detected. Cameras were mounted 30–40 cm above ground. Thirty cameras were deployed in each array, yielding a total sampling effort of 6,900 camera-trap days in the eastern sector and 4,740 camera-trap days in the western sector. Due to camera malfunctions, data from five cameras in the western survey were excluded from analyses, resulting in 25 functional cameras contributing to the final dataset.

**Figure 2:**
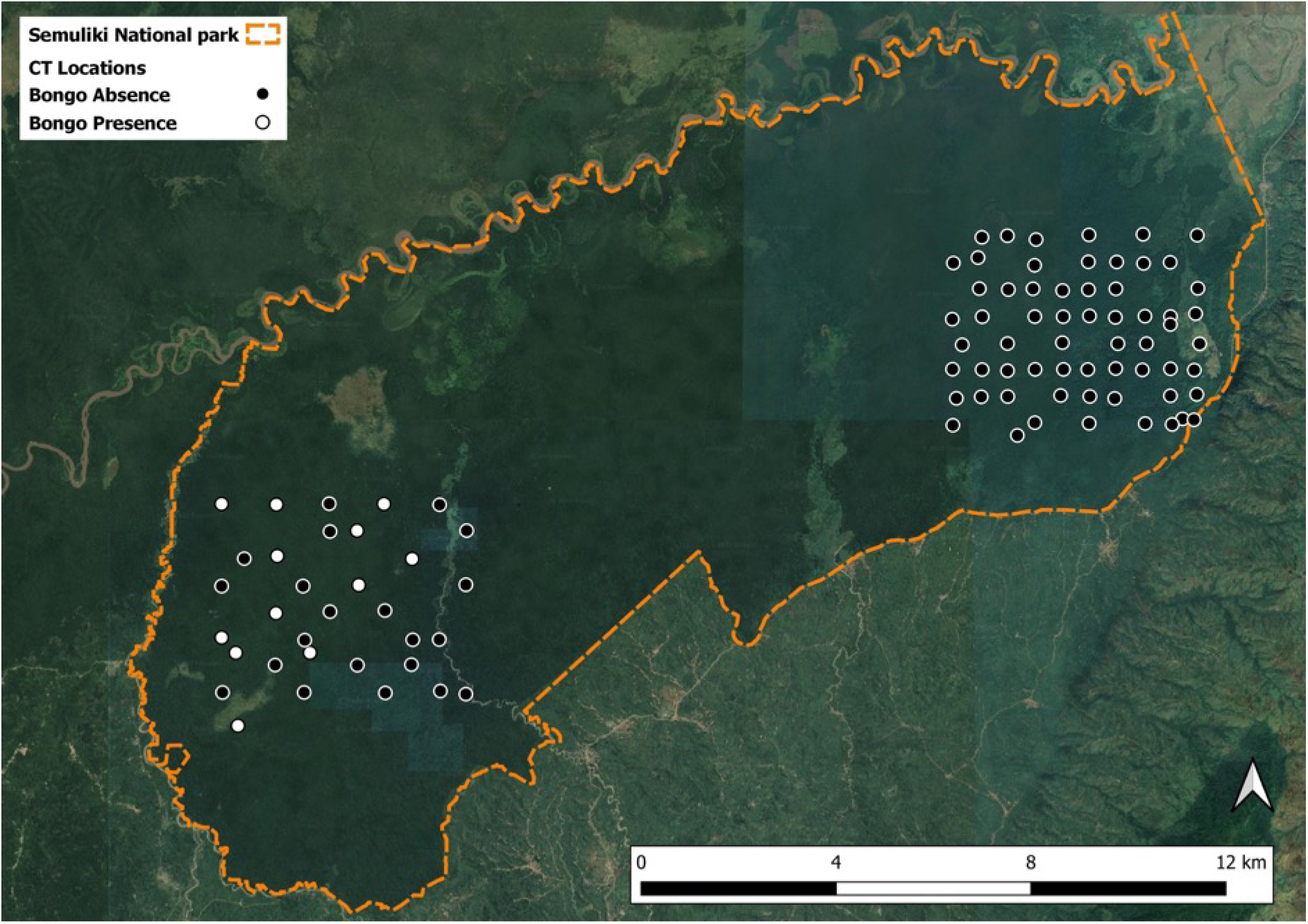
Location of camera trap surveys conducted in Semuliki National Park between October 2019 and August 2020. Two consecutive survey arrays were deployed in the eastern sector (October–December 2019 and December 2019–January 2020), followed by one survey array in the western sector (March–August 2020). Western bongo were detected only during the western-sector survey; camera trap locations with bongo detections are shown in white. Camera trap locations in the eastern sector were not treated as confirmed absences because seasonal variation in habitat use and detectability could not be excluded.

### Individual Bongo Identification

Individual bongos were identified based on unique flank stripe patterns (Sandri et al., 2023). Capture histories were constructed by treating individuals as “marked” at first detection and subsequent detections as recaptures. Due to asymmetry in flank patterns, left and right flanks were initially treated as independent capture units, and separate capture histories were compiled for each flank type to avoid double counting (Sandri et al., 2023; Koopmans et al., 2021). For individuals where left and right flanks could be reliably matched, detections were linked across sides and combined into individual capture histories, increasing the number of available recaptures for analysis while avoiding duplicate counts of the same individual.

### Habitat Covariates

We derived environmental covariates from Landsat 8 imagery (OLI/TIRS; scene LC81730592020134LGN00; U.S. Geological Survey, 2020). Tasseled Cap Transformation (TCT; Baig et al., 2014) was applied to extract three orthogonal indices: brightness (TCB), greenness (TCG), and wetness (TCW). These indices capture variation in vegetation structure, productivity, and moisture conditions respectively. Vegetation structure was further characterized using forest canopy height from the GLAD/UMD Global Forest Canopy Height dataset (Potapov et al., 2020). Anthropogenic disturbance was represented by Euclidean distance to villages and roads, calculated in QGIS (QGIS Development Team, 2022). Terrain variables included distance to rivers, generated from the Waterbodies in Uganda GIS dataset distributed by the World Resources Institute (WRI), and terrain ruggedness, derived from a 30 m Shuttle Radar Topography Mission (SRTM) digital elevation model (Farr et al., 2007).

All covariates were aggregated to 100 m using the median value (package *raster*; Hijmans, 2025) to better reflect the spatial scale at which bongo likely respond to habitat (Estes et al., 2011). Collinearity among predictors was assessed using variance inflation factor (Cavada et al., 2017), with a threshold of 3. Variables retained for modelling were canopy height, distance to villages, ruggedness, TCG, and TCW.

Prior to modeling, all spatial covariates were standardized (mean = 0, SD = 1) to facilitate model convergence and interpretability of coefficients. Extreme values that could disproportionately influence density predictions were winsorized to the 1st and 99th percentiles to reduce the effect of outliers while retaining overall variation (Zuur et al., 2009).

### Spatially Explicit Capture Recapture model

Because bongos were detected only in the western-sector survey, analyses were restricted to this dataset (25 cameras for 3233 camera-trap days). We used a half-normal detection function, where detection probability is defined by a baseline encounter probability (g₀) and a spatial scale parameter (σ), which describes the decline in detection probability with distance to the center of activity (Efford, 2025).

Capture histories were constructed using 7-day sampling occasions, resulting in 17 occasions across the survey period between March and August 2020. The spatial extent of the model was defined by a buffer surrounding camera trap locations in the western sector survey, calculated in the R package *secr* (Efford, 2025) using 4 × σ estimated via the function *RPSV*. This resulted in a buffer of 4,030.5 m. The resulting state-space covered 140 km^2^.

To identify the factors influencing bongo density, we adopted a two-stage modelling approach. In the first stage, we fitted seven candidate spatially explicit capture–recapture (SECR) models each including a single habitat covariate as a predictor of density (D) plus a null model containing no habitat covariate. Given the limited number of detections and identified individuals, models containing multiple habitat covariates or interactions were not considered, as they would have led to overparameterization and unsupported complexity (Burnham & Anderson, 2002). Candidate models were compared using an information-theoretic approach based on Akaike’s Information Criterion corrected for small sample size (AICc; Burnham & Anderson, 2002), and the best-supported habitat model was retained. In the second stage, we incorporated sex as an additional covariate affecting the relevant model parameters, generating eight candidate models based on the best-supported habitat model. These models were again ranked using AICc to identify the final model. We assessed the precision of density estimates using the coefficient of variation of density (CVD; Efford & Fewster, 2013). Abundance was estimated using the *region. N* function in the secr package in R.

## RESULTS

Bongos were recorded in 17 independent detection events, defined as consecutive captures separated by at least 20 minutes, across nine camera-trap stations. A total of 32 flanks were identified, comprising 18 right and 14 left flanks. These included 22 flanks (11 right, 11 left) of females, of which five were from immature individuals, and 10 flanks (7 right, 3 left) of males, of which two were from immatures. Using matched left and right flanks, we identified 10 individuals: seven females (three adults and four immatures) and three males (two adults and one immature). These individuals were combined with the right-flank capture histories, which provided the larger sample, resulting in a final dataset of 18 identified individuals for SECR analyses, with an observed sex ratio of 0.64 males per female. Although matching left and right flanks did not increase the number of identified individuals, it increased the number of recapture events from 8 to 18.

The candidate SECR models are summarised in Table 1. In the first stage of model selection, which evaluated habitat covariates, the best-supported model included only TCG as a predictor of density (Table 1A). In the second stage, sex was added as a covariate to this model. The best-supported overall model included both TCG and sex as predictors of density (Table 1B). Model selection was based on AICc, with preference given to models receiving strong support (ΔAICc ≤ 2) and acceptable precision (CVD < 0.30). The selected model had a CVD of 0.29, indicating moderate precision.

**Table 1:**
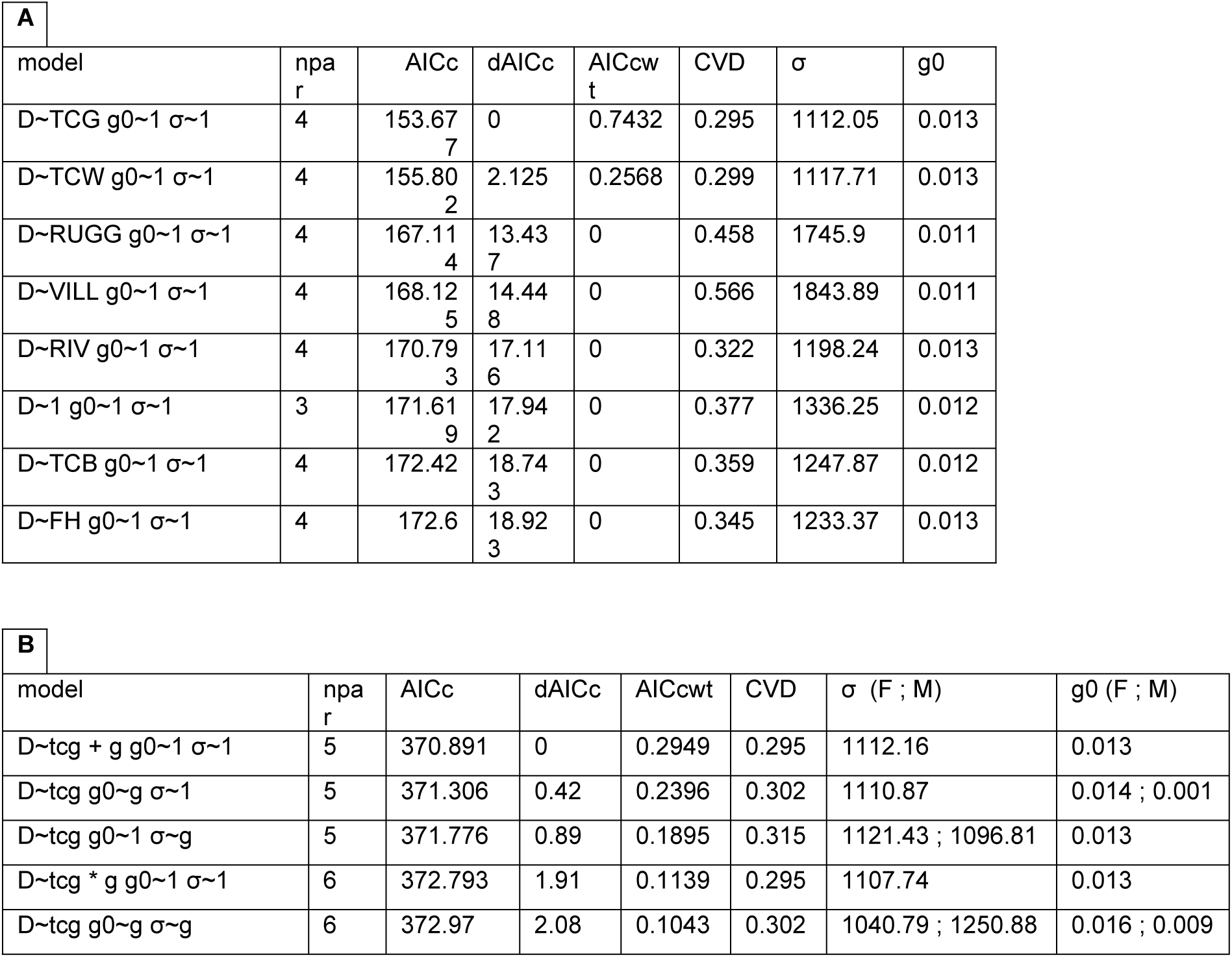
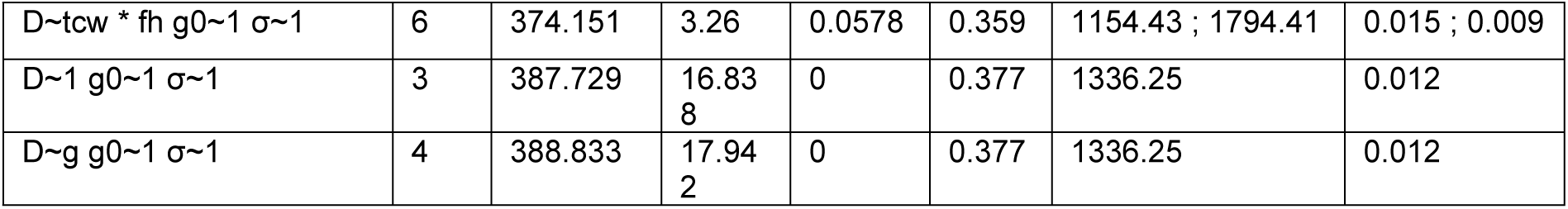
Model selection results for the two-stage spatially explicit capture–recapture (SECR) analysis of bongo density in Semuliki National Park. (A) Candidate models evaluating the influence of individual habitat covariates on density (D). Each model included a single habitat covariate, while baseline detection probability (g₀) and the spatial scale parameter (σ) were held constant. (B) Candidate models evaluating the influence of sex (g) on the best-supported habitat model identified in stage one (TCG). Sex was fitted as a covariate on density (D), baseline detection probability (g₀), and/or the spatial scale parameter (σ), with competing models ranked using AICc. Models are ranked by Akaike’s Information Criterion corrected for small sample size (AICc). Reported are the number of estimated parameters (npar), AICc, ΔAICc, Akaike weight (AICcwt), coefficient of variation of the density estimate (CVD), estimated movement parameter (σ), and baseline detection probability (g₀). Single values indicate no sex effect; paired values are female and male estimates, respectively.

The top-ranked model (D ~ TCG + sex, g0 ~ 1, σ ~ 1) estimated a density of 0.58 individuals km^−2^ (SE = 0.171; 95% CI = 0.32–1.09). Extrapolation of the fitted model across the full extent of Semuliki National Park yielded an estimated abundance of 30.40 females (SE = 8.83; 95% CI = 13.08–47.71) and 25.89 males (SE = 6.53; 95% CI = 13.09–38.69), corresponding to a total population estimate of 56.29 individuals (SE = 10.99; 95% CI = 34.76–77.82) with a model-estimated sex ratio of 0.85 males per female. These park-wide estimates were similar to those obtained within the 140 km^2^ fitted state space (56.49 individuals; 30.53 females and 25.97 males), indicating that areas added during extrapolation contributed little to the predicted abundance under the fitted TCG–density relationship.

## DISCUSSION

This study provides the first estimates of density and abundance for the only known population of Western bongo in East Africa, located in Semuliki, Uganda. Density estimates were associated with habitat variables reflecting forest structure (TCG) consistent with the species’ ecology as a forest habitat specialist. The estimated density of 0.58 individuals/km^2^ falls within the range of published values for the species. Previous estimates vary widely, from 0.08 individuals/km^2^ in a hunting concession in the Republic of Congo (Koopmans et al., 2021) to 0.25–1.2 individuals/km^2^ in South Sudan and Dzangha Sangha - Central African Republic (Hillman, 1986; Klaus-Hugi et al., 1999), and up to 5.3 individuals/km^2^ in Kakum National Park, Ghana (Kwaku et al., 2014). Estimates for the critically endangered Eastern bongo in Kenya are lower, ranging from 0.03 to 0.07 individuals/km^2^ (Sandri et al., 2023). The variation among published estimates likely reflects differences in both ecological context and analytical approaches. Populations occurring in areas with contrasting habitat quality, protection levels, and hunting pressure may naturally differ in density, while methodological differences—including survey design, individual identification methods, and whether spatial variation in habitat suitability was incorporated—can further influence estimates. The use of SECR models with habitat covariates in this study allowed density to vary spatially, potentially providing a more representative estimate for a heterogeneous landscape such as Semuliki.

Sex ratio among identified individuals was less skewed than reported in other bongo populations (e.g. Hillman, 1983; Sandri, Prettejohn et al., 2023). While this may indicate a relatively balanced population structure, it may also reflect sex-specific differences in movement and detectability. The SECR model estimated a less biased sex ratio (0.85 males per female) compared with the observed ratio among identified individuals (0.64 males per female), suggesting that part of the observed female bias may result from differences in detection rather than true population structure. Male bongos are typically more solitary and mobile, whereas females are more likely to form stable groups (Sandri et al., 2023), potentially reducing the likelihood of male detection. More broadly, this highlights that demographic patterns inferred from field observations can reflect both underlying population processes and the way individuals are observed, particularly when sexes differ in behaviour, movement, or detectability (Yoccoz et al., 2001; Anderson, 2001). The presence of immature individuals suggests reproduction within the park, supporting the interpretation that at least part of the population is established within Semuliki, although further data are required to confirm patterns of residency and movement.

Detections were concentrated within the western sector of the park, suggesting that the population may be restricted to a relatively limited area. This pattern is consistent with the species’ association with specific habitat conditions (Estes et al., 2010; Sandri 2020) and is supported by the absence of detections in repeated surveys in the eastern sector. However, the potential influence of low detection probability and seasonal movements cannot be excluded. The habitat-based SECR model predicted that additional areas beyond the surveyed state space contributed little to overall abundance, suggesting that bongo distribution within Semuliki may be spatially restricted. The restricted contribution of areas outside the surveyed area to predicted abundance suggests that bongo distribution within Semuliki may be associated with the spatial distribution of suitable habitat, as represented by TCG. This provides some support for the possibility that the absence of detections during eastern-sector surveys reflected genuinely low density rather than solely imperfect detection. However, because TCG represents a static landscape characteristic, this inference does not account for potential seasonal shifts in habitat use or movements. Seasonal variation in resource availability or ranging behaviour, could result in temporary absence from areas that remain suitable based on habitat characteristics alone. As a baseline assessment, this study provides an initial understanding of bongo density, abundance, and distribution within Semuliki. Future monitoring across seasons will be essential to determine whether the observed spatial patterns represent persistent habitat associations or reflect seasonal variation in habitat use, thereby improving the basis for long-term conservation planning.

Habitat associations alone may not fully explain the observed distribution pattern. Anthropogenic pressure may also contribute to this spatial pattern. Earlier camera trap surveys conducted in 2018 and in the present study recorded evidence of human activity at multiple locations, including bushmeat hunting and resource extraction, although overall levels were relatively low. Signs of hunting activity, including the use of snares targeting medium to large-bodied ungulates, were recorded in areas away from established patrol routes, indicating spatial variation in protection effectiveness (Nixon, pers. Comm.). Such pressures may influence habitat use and local distribution of bongos as hunting and human disturbance can alter the distribution and behaviour of large forest ungulates (Abernethy et al., 2013). Semuliki has also experienced a complex history of land use and protection, with periods of limited enforcement and human settlement (Kingdon 1971), it was designated as a forest reserve from 1932 to 1993 when it became a national park (Chege et al., 2002). These historical processes may have contributed to variation in forest structure across the park, with potential implications for species distributions over time. This is particularly relevant for bongo, as the habitat associations identified in this study suggest that variation in forest structure may contribute to their current distribution within Semuliki. Previous studies have shown that bongos can associate with areas of secondary forest and dense vegetation, highlighting the importance of forest structure in shaping habitat suitability for the species (Estes et al., 2008; Sandri, 2020).

The transboundary context of Semuliki, contiguous with Virunga National Park in the Democratic Republic of Congo, places this bongo population within the wider Bwamba forest landscape described by Kingdon (1971) and the broader Albertine Rift biodiversity hotspot (Plumptre et al., 2007). Although forest habitats extend across political boundaries, differences in governance, protection capacity, and human activity between protected areas may influence patterns of hunting pressure and wildlife movement (Abernethy et al., 2013). Together, historical disturbance, ongoing human activity, and spatial variation in protection may have contributed to the current distribution of bongos within Semuliki.

Knowledge of Western bongo status across its range remains limited, and few studies have produced robust density estimates. This is of concern given ongoing pressures from hunting and habitat disturbance, and the lack of comprehensive regulation across parts of the species’ range (Koopmans et al., 2021). The discovery and assessment of this population in Semuliki therefore represents an important opportunity to establish long-term monitoring and conservation efforts. Opportunistic field observations during camera deployment, including tracks, dung, and limited direct sightings, further support the presence of a small and elusive population within the park (Nixon, pers. comm). As a large-bodied, slow-reproducing forest ungulate, bongos are likely to be particularly sensitive to hunting pressure, including indiscriminate snaring. This reinforces the importance of effective protection measures, particularly in areas where human activity occurs beyond regular patrol coverage.

This study is subject to several limitations common to low-density, logistically constrained surveys of rare species. Individual identification relied primarily on single-flank photographs; discarding or restricting partial-identity capture histories in this way can reduce precision and introduce bias into abundance estimates (Augustine et al. 2018). Full implementation of spatially explicit partial-identity models was beyond the scope of the present dataset but should be prioritised as data accumulate. Female bongos are known to associate in loose groups, which may violate the assumption of independent detection underlying SECR. Simulation studies suggest that low-to-moderate spatial aggregation and behavioural cohesion have limited effect on the bias of density estimates, though they can inflate overdispersion and reduce the reliability of reported confidence intervals (Bischof et al. 2020). Our estimates of precision should be interpreted with this in mind. The relatively small number of detections and identified individuals limited the complexity of models that could be robustly supported: the cumulative discovery curve of identified individuals had not reached a clear asymptote by the end of the survey, a pattern also reported in other SECR studies of low-density, wide-ranging species (e.g., Maputla et al. 2013), and finite mixture approaches to unmodelled individual heterogeneity were not pursued as they require substantially larger sample sizes than obtained here to yield reliable, comparable estimates (Marrotte et al. 2022). Sex was retained as the most defensible, biologically meaningful covariate available given these constraints. Habitat covariates were similarly constrained by the resolution and availability of remotely sensed data, and the relationships identified should be interpreted as indicative rather than definitive. Increased sampling effort, spatial replication, and long-term monitoring will be important for refining these estimates, improving understanding of bongo ecology in the region, and determining how population patterns vary through time. These efforts will also be essential for translating baseline assessments such as this into effective long-term conservation strategies.

This study provides an initial baseline of bongo density, abundance, and distribution in Semuliki National Park, establishing a foundation for future population monitoring and conservation planning. The apparent concentration of individuals within a restricted portion of the park may have important conservation implications, as local populations occupying limited areas may be disproportionately affected by site-specific disturbances, even when the species as a whole has a broader distribution (Hanski, 1998). However, continued monitoring across seasons will be essential to determine whether the observed spatial patterns reflect persistent habitat associations or temporal shifts in distribution. Such long-term monitoring will be critical for refining our understanding of bongo population dynamics and ensuring that conservation actions are targeted effectively. Protecting the bongo across its range will ultimately depend on understanding and conserving the individual populations that persist in forests such as Semuliki; by improving our understanding of this one remaining population at the edge of its distribution, this study contributes to the broader effort to conserve Africa’s largest forest antelope.

## AUTHOR CONTRIBUTION

Conceptualization: SN, NM, SI, MV, RA, TS

Field Work: SN, NM, SI, MV

Data curation: NM, TS, AvH

Formal analysis: TS, AvH

Writing – original draft: TS

Writing – review & editing: All Authors

